# The mitochondrial Ca^2+^ uniporter MCU is required for normal glucose-stimulated insulin secretion *in vitro* and *in vivo*

**DOI:** 10.1101/781161

**Authors:** Elizabeth Haythorne, Eleni Georgiadou, Matthew T. Dickerson, Livia Lopez-Noriega, Timothy J. Pullen, Gabriela da Silva Xavier, Samuel P.X. Davis, Aida Martinez-Sanchez, Francesca Semplici, Rosario Rizzuto, James A. McGinty, Paul M. French, Matthew C. Cane, David A. Jacobson, Isabelle Leclerc, Guy A. Rutter

## Abstract

Mitochondrial oxidative metabolism is central to glucose-stimulated insulin secretion (GSIS). Whether Ca^2+^ uptake into pancreatic β-cell mitochondria potentiates or antagonises this process is still a matter of debate. Although the mitochondrial importer (MCU) complex is thought to represent the main route for Ca^2+^ transport across the inner mitochondrial membrane, its role in β-cells has not previously been examined *in vivo*. Here, we inactivated the pore-forming subunit MCUa (MCU) selectively in the β-cell in mice using *Ins1*Cre-mediated recombination. Glucose-stimulated mitochondrial Ca^2+^ accumulation, ATP production and insulin secretion were strongly (p<0.05 and p<0.01) inhibited in *MCU* null animals (βMCU-KO) *in vitro*. Interestingly, cytosolic Ca^2+^ concentrations increased (p<0.001) whereas mitochondrial membrane depolarisation improved in βMCU-KO animals. Male βMCU-KO mice displayed impaired *in vivo* insulin secretion at 5 (p<0.001) but not 15 min. post intraperitoneal (IP) injection of glucose while the opposite phenomenon was observed following an oral gavage at 5 min. Unexpectedly, glucose tolerance was improved (p<0.05) in young βMCU-KO (<12 weeks), but not older animals. We conclude that MCU is crucial for mitochondrial Ca^2+^ uptake in pancreatic β-cells and is required for normal GSIS. The apparent compensatory mechanisms which maintain glucose tolerance in βMCU-KO mice remain to be established.

## Introduction

Defective insulin secretion underlies diabetes mellitus, a disease affecting almost 1 in 8 of the adult population worldwide (https://www.idf.org/). The most prevalent form of this condition is Type 2 diabetes (T2D) where pancreatic β-cell failure usually, though not always, occurs in the face of insulin resistance in other tissues (1). Current therapeutic strategies have limited efficacy and there remains a desperate need to develop more effective treatments to tackle this growing epidemic.

Pancreatic β-cells ensure blood glucose homeostasis by responding to a rise in circulating nutrient levels with insulin secretion. Glucose-induced increases in mitochondrial oxidative metabolism are central to the stimulation of insulin secretion, and drive an increase in cytosolic ATP:ADP ratio, closure of ATP-sensitive K^+^ (K_ATP_) channels, Ca^2+^ influx and exocytosis (2). Ca^2+^ ions are also taken up by mitochondria (3; 4) and this has been suggested to activate tricarboxylate (TCA) cycle and other intra-mitochondrial enzymes (5) in order to enhance the production of reducing equivalents for the electron transport chain and ATP generation (2). Although a number of approaches have been used previously to explore the role of intra-mitochondrial Ca^2+^ in controlling insulin secretion, the role of these ions in modifying ATP synthesis, and hence exocytosis, in these cells is still debated (6; 7).

Importantly, there is accumulating evidence to suggest that mitochondrial dysfunction in the pancreatic β-cell leads to impaired glucose-stimulated insulin secretion and may contribute to the development of T2D (8). Moreover, glucolipotoxicity impairs mitochondrial Ca^2+^ uptake in isolated β-cells (9), suggesting a possible role in defective secretion under these conditions.

The mitochondrial Ca^2+^ uniporter (MCU) forms the Ca^2+^-selective pore of a multiprotein MCU-complex including MCUa [MCU], MICU1-3, MICUR1, and EMRE which allows Ca^2+^ entry into mitochondria (10). *In vitro* and *in vivo* models of MCU silencing or ablation have revealed a robust reduction in mitochondrial Ca^2+^ uptake associated with blunted Ca^2+^-dependent activation of the TCA cycle, oxygen consumption, ATP production (9; 11; 12) and mitochondrial reactive oxygen species generation (13). Whole body *Mcu* knockout (KO) mice display normal basal cardiac parameters even though mitochondria isolated from cardiac myocytes display impaired Ca^2 +^ uptake and Ca^2 +^-dependent oxygen consumption. Interestingly, resting ATP levels are unaltered in muscle cells in *Mcu*-KO mice, suggesting that MCU depletion does not affect basal mitochondrial metabolism. *Mcu*-KO mice similarly display only reduced maximal muscle power in association with reduced metabolic flux and activity of TCA cyle enzymes in skeletal muscle mitochondria (11). Given that both cardiac and skeletal muscles are highly metabolically active tissues, it is surprising that whole body *Mcu*-KO mice display a mild phenotype (11–13). However, glucose homeostasis and insulin secretion were not examined in detail in these earlier studies.

We have previously shown that reducing glucose-stimulated mitochondrial Ca^2+^ uptake in rodent pancreatic β-cells through knockdown of *MCU in vitro* impairs the sustained increase in ATP:ADP ratio usually seen in response to high glucose and ablates sulfonylurea-stimulated insulin secretion (9; 14). Similar findings were also made in clonal β-cells (15). However, these earlier studies provided no insights as to the impact of reducing mitochondrial Ca^2+^ uptake on glucose-stimulated insulin secretion *in vivo*, nor how this may, in turn, impact the physiology of the living animal.

Tissue-specific manipulation of MCU activity provides an alternative and powerful means to examine the role of mitochondrial Ca^2+^ in particular tissue or cell type. In the present study we have therefore generated mice in which *Mcu* is deleted highly selectively in the pancreatic β-cell, and explored the impact on insulin secretion and whole body glucose homeostasis. We show that mitochondrial Ca^2+^ uptake, glucose-induced ATP increases and insulin secretion are substantially impaired *in vitro* in dissociated or islets from KO mice. Insulin release is also impaired in the living mouse, despite improvements in glucose tolerance.

## Materials and Methods

### Generation of β-cell specific MCU-KO mice

All *in vivo* procedures were approved by the UK Home Office according to the Animals (Scientific Procedures) Act 1986. Animals were fed *ad libitum* with a standard mouse chow diet (Research Diets, Inc) unless otherwise stated. For high fat/sucrose diet (HFHS) treatment, mice were placed on diet at 5-6 weeks of age for 2 weeks (58% [wt/wt] fat and sucrose content; Research Diets, Inc) prior to analysis.

C57BL/6J mice bearing *Mcu* (also termed *Ccdc109a* and *C10orf42*) alleles with *Flox*P sites flanking exons 11 and 12 were generated by GenOway (Grenoble, Fr) and bred to animals carrying *Cre* recombinase inserted at the *Ins1* locus (*Ins1*Cre) (16). Use of this Cre line allowed efficient and β-cell-selective deletion of both *Mcu* splice variants (βMCU-KO), without recombination in the brain or confounding expression of human growth hormone. Mice bearing *floxed Mcu* alleles but lacking *Cre* recombinase were used as littermate controls (WT). Possession of *Ins1*Cre alleles alone exerted no effect on glycaemic phenotype (not shown). The sequences of primers used for genotyping and qRT-PCR for *Mcu*, *Kcnj11* and *Abcc8* are provided under Supplemental Tables 1 and 2.

For measurements of mRNA levels, pancreatic islets were isolated by collagenase digestion (17). Deletion of *Mcu* was determined using quantitative real-time PCR. RNA was extracted from islets using Trizol (Invitrogen) and reverse transcribed using a high capacity reverse transcription kit (Invitrogen) (18). Relative gene expression was determined using SYBR Green (Invitrogen), and expression of *Mcu* was normalised to β-actin mRNA.

### Intraperitoneal glucose, insulin tolerance tests and measurement of insulin secretion in vivo

To investigate glucose tolerance, male or female mice (ages 8-24 weeks as indicated) were fasted overnight for 16 h before injection of glucose solution (20% w/v, 1g/kg body weight) intraperitoneally. Glucose was measured in tail vein blood at 0, 5, 15, 30, 60, 90 and 120 min. using an ACCU-CHECK Aviva glucometer (Roche) (19).

To ascertain insulin tolerance, mice were fasted for 5 h before human insulin (0.75 units/kg body weight, Sigma Aldrich) was injected intraperitoneally. Blood glucose was measured in tail vein blood at 0, 15, 30 and 60 min. (19).

For *in vivo* insulin secretion experiments, animals were fasted overnight for 16 h and glucose (20% w/v, 3 g/kg body weight) was either given intraperitoneally or oral gavage. Plasma was separated by centrifugation and insulin was measured using an ultra-sensitive mouse insulin ELISA kit (CrystalChem).

### Single cell fluorescence imaging

Pancreatic islets were isolated as above, dissociated into single β-cells and plated onto glass coverslips (9; 20). Mitochondrial Ca^2+^ uptake was measured via adenovirus-mediated delivery of a mitochondrially-targeted recombinant Ca^2+^ probe, R-GECO (21). Cells were infected and incubated for 48 h prior to imaging in Krebs-Ringer bicarbonate buffer (140 mM NaCl, 3.6 mM KCl, 0.5 mM NaH_2_PO_4_, 2 mM NaHCO_3_ (saturated with CO_2_), 1.5 mM CaCl_2_, 0.5 mM MgSO_4_, 10 mM HEPES and 3 mM D-glucose, pH7.4). To examine ATP:ADP changes in response to a rise in extracellular glucose concentration, dissociated β-cells were infected with an adenovirus bearing cDNA encoding the ATP sensor Perceval (9) and incubated for 48 h prior to fluorescence imaging. In all experiments, cells were equilibrated for at least 30 min. in Krebs-Ringer bicarbonate containing 3 mM glucose prior to the start of acquisitions. Excitation/emission wavelengths were (nm): 490/630 and 410/630 (Fura-Red), 530/590 (R-GECO) and 470/535 (Perceval). All imaging experiments were performed using an Olympus IX71 microscope with 40x magnification objective, an F-View-II camera and an MT-20 excitation system equipped with a Hg/Xe arc lamp with image capture at 0.2 Hz excitation time 50 ms).

For experiments with tetramethylrhodamine (TMRE), β-cells were loaded with 10nM TMRE in imaging buffer (140 mM NaCl, 3.6 mM KCl, 0.5 mM NaH_2_PO_4_, 24 mM NaHCO_3_ (saturated with CO_2_), 1.5 mM CaCl_2_, 0.5 mM MgSO_4_, 10 mM HEPES and 3 mM glucose) for 45 min. and re-equilibrated with 2nM TMRE for 10 min. before recordings. TMRE (2nM) was present throughout, and cells excited at 550 nm. FCCP (Carbonyl cyanide-4-phenylhydrazone, 1μM) was administrated as indicated and imaging performed using a Zeiss AxioObserver microscope using 63x 1.4NA oil objective, a Hamamatsu Flash 4 camera and an LED (light emitting diode) excitation system (excitation filter 534/20nm and emission filter 572/28nm) at 0.3 Hz (250 ms exposure). Traces represent mean normalised fluorescence intensity (F/F_min_) over time, where F_min_ is the average fluorescence recorded at 3mM glucose.

### Whole islet fluorescence imaging

Ca^2+^ imaging of whole islets was performed after loading with Cal-520 AM (Stratech;2 μM), or mito Pericam adenovirus (MOI 100), and perifusion in Krebs-Ringer bicarbonate buffer containing 3mM or 17mM glucose, 17mM glucose with 0.1mM diazoxide or 20mM KCl. Images were captured at 0.5Hz on a Zeiss Axiovert microscope equipped with a 10X 0.3–0.5 NA oil immersion objective, coupled to a Nipkow spinning-disk head (Yokogawa CSU-10) and illuminated at 491nm. Data were analysed using ImageJ with a purpose-designed macro (available upon request).

### In vitro insulin secretion

Insulin secretion assays were performed on islets isolated from male mice (8-10 weeks) (24). Secreted and total insulin were quantified using a homogeneous time-resolved fluorescence (HTRF) insulin kit (Cisbio) in a PHERAstar reader (BMG Labtech, UK) following the manufacturer’s guidelines. Data are presented as secreted insulin/insulin content.

### Electrophysiology

Electrophysiological recordings were performed on single β-cells in perforated patch-clamp configuration using an EPC9 patch-clamp amplifier controlled by Pulse acquisition software (Heka Elektronik, Pfalz, Germany). β-cells were identified morphologically and by depolarisation of the membrane potential in response to 17 mM glucose. β-cells were constantly perfused at 32 °C with normal saline solution (135mM NaCl, 5mM KCl, 1mM MgCl_2_, 1mM CaCl_2_, 10mM HEPES). Recording electrodes had resistances of 8-10 MΩ and were filled with a solution comprised of 140mM KCl, 5mM MgCl_2_, 3.8mM CaCl_2_, 10mM HEPES, 10mM EGTA (pH 7.2) and 20–25 μg/ml amphotericin B (Sigma-Aldrich).

Voltage-dependent calcium channel (VDCCs) currents were recorded from dispersed mouse β-cells as previously described (22).

### β-cell mass

Whole pancreatic optical projection tomography, to 19 µm resolution, was performed as described (23).

### Statistical Analysis

Data are expressed as mean ± SEM except for small datasets (n<3), where data are shown as mean ± S.D. Significance was tested by Student’s two-tailed t-test, one or two-way ANOVA with Sidak’s or Bonferroni multiple comparison test for comparison of more than two groups, using GraphPad Prism 7 software. *p*<0.05 was considered significant.

## Results

### *Mcu* ablation from pancreatic β-cells attenuates GSIS *in vitr*o

To ablate MCU from pancreatic β-cells, animals bearing alleles in which *Lox*P sites flanked exons 11 and 12 of the *Mcu* gene were generated (See Materials and Methods) and crossed to mice harbouring *Cre* recombinase inserted at the *Ins1* locus (16) (Fig. 1A). This strategy ensured targeted deletion of the longer splice variant of *Mcu*, which predominates in islets (results not shown), whilst excluding any compensatory increase in the expression of the shorter variant. *Mcu* deletion was confirmed by qRT-PCR (Fig. 1B). Relative to β-Actin (*Actb*), expression of the *Mcu* transcript in KO islets was decreased by ∼75% *vs* control islets (p<0.05; Fig. 1B). This level of reduction is consistent with near-complete elimination of *Mcu* mRNA from β-cells, assuming a β-:α-cell ratio of ∼3:1 (24) and similar levels of *Mcu* expression in each cell type in WT islets (25).

**Figure 1.**
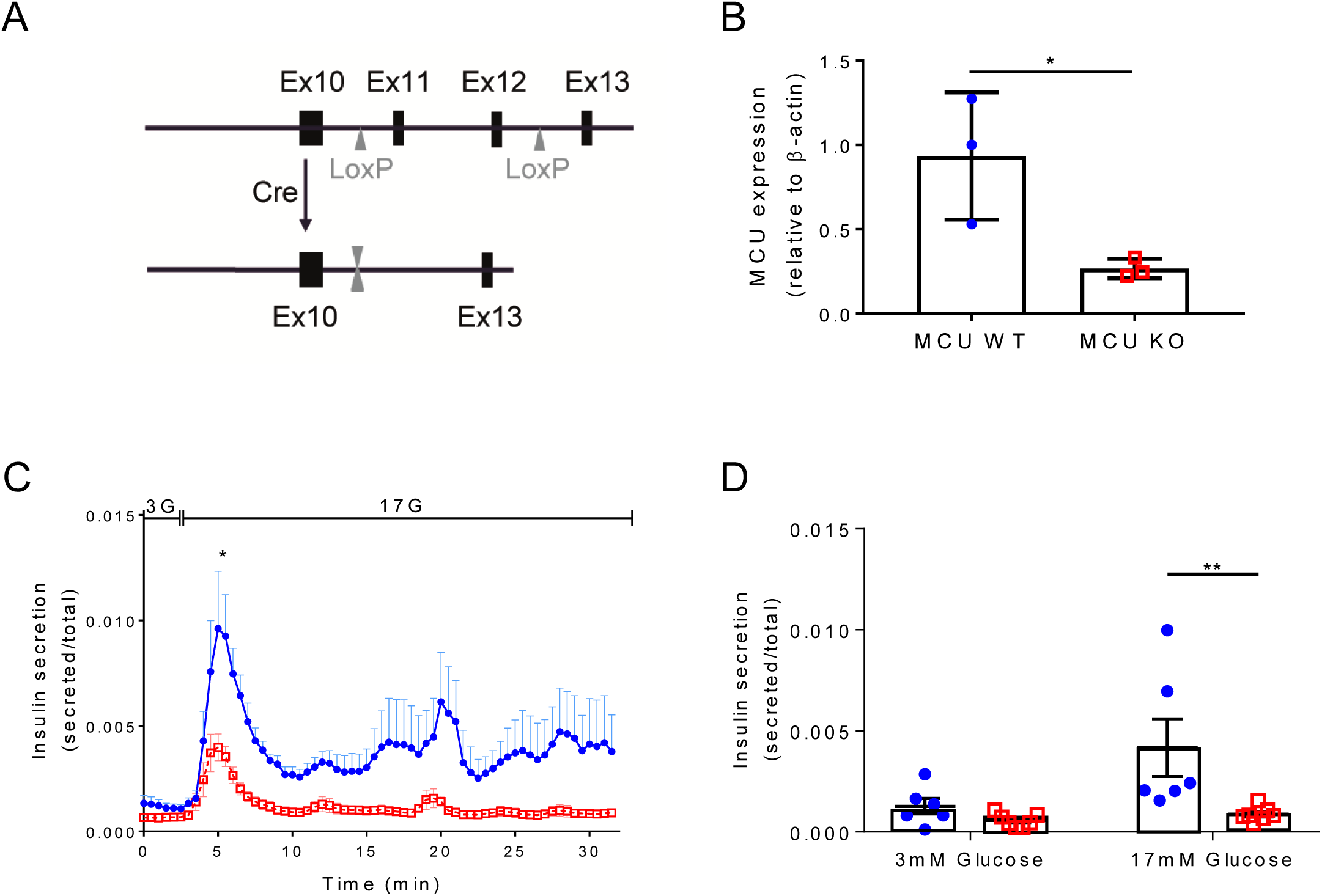
Isolated islets from βMCU-KO mice display attenuated glucose-stimulated insulin secretion *in vitro*. (A) Gene deletion was achieved by breeding mice carrying *Mcu* alleles with FloxP sites flanking exons 11 and 12 with mice bearing Cre recombinase inserted at the *Ins1* locus. (B) qRT-PCR quantification of *Mcu* expression (*p<0.05; *n=*3 mice per genotype). Values represent mean ± SD. (C) Insulin secretion from islets isolated from βMCU-WT and KO mice during perifusion and (D) serial incubations of islets in batches at 3 mM (3G) or 17 mM glucose (17G). A significant decrease in insulin secretion was observed in islets isolated from KO mice during the first peak (A; 4-8 min., p<0.05, *n=*4-5 mice per genotype; unpaired two-tailed Student’s t-test). In (D), a significant decrease was observed between genotypes in secretion stimulated by 17 mM glucose (p<0.01; two-way ANOVA test and Sidak’s multiple comparisons test) compared to WT mice (*n=*6 WT, *n=*7 KO animals). Blue circles, βMCU-WT, red squares, βMCU-KO mice.

We next explored the consequences for glucose-stimulated insulin secretion in isolated βMCU-KO islets. In perifusion experiments, βMCU-KO islets displayed a significant blunting in the secretory response to elevated glucose, with the attenuation in insulin release most evident at high glucose concentrations (17 mM), as determined at the first peak (p<0.05; Fig. 1C). These results were confirmed by independent experiments involving batch incubation of islets (p<0.01; Fig. 1D). In contrast, βMCU-KO islets displayed no difference *vs* control islets in insulin secretion stimulated by depolarisation with 20 mM KCl in either system (data not shown).

### MCU deletion from pancreatic β-cells impairs glucose-stimulated mitochondrial but not cytosolic Ca^2+^ uptake

Since MCU provides the main route for Ca^2+^ entry into mitochondria in other cell types (26), we first determined the impact of deleting *MCU* on this process in pancreatic β-cells. Changes in mitochondrial free Ca^2+^ concentration ([Ca^2+^]_mt_) were investigated by live cell fluorescence microscopy using whole or dissociated islets expressing an adenovirus (mito Pericam), or a genetically-encoded Ca^2+^ indicator (R-GECO), respectively (21). After pre-incubation in the presence of low (3 mM) glucose, increases in [Ca^2+^]_mt_ were provoked in control islets or β-cells by stimulation with high (17 mM) glucose or plasma membrane depolarisation with KCl (20 mM). A depolarising K^+^ concentration and the K_ATP_ channel opener diazoxide were then deployed together to bypass glucose regulation of the K_ATP_ channel (Fig. 2C’, E’).

**Figure 2.**
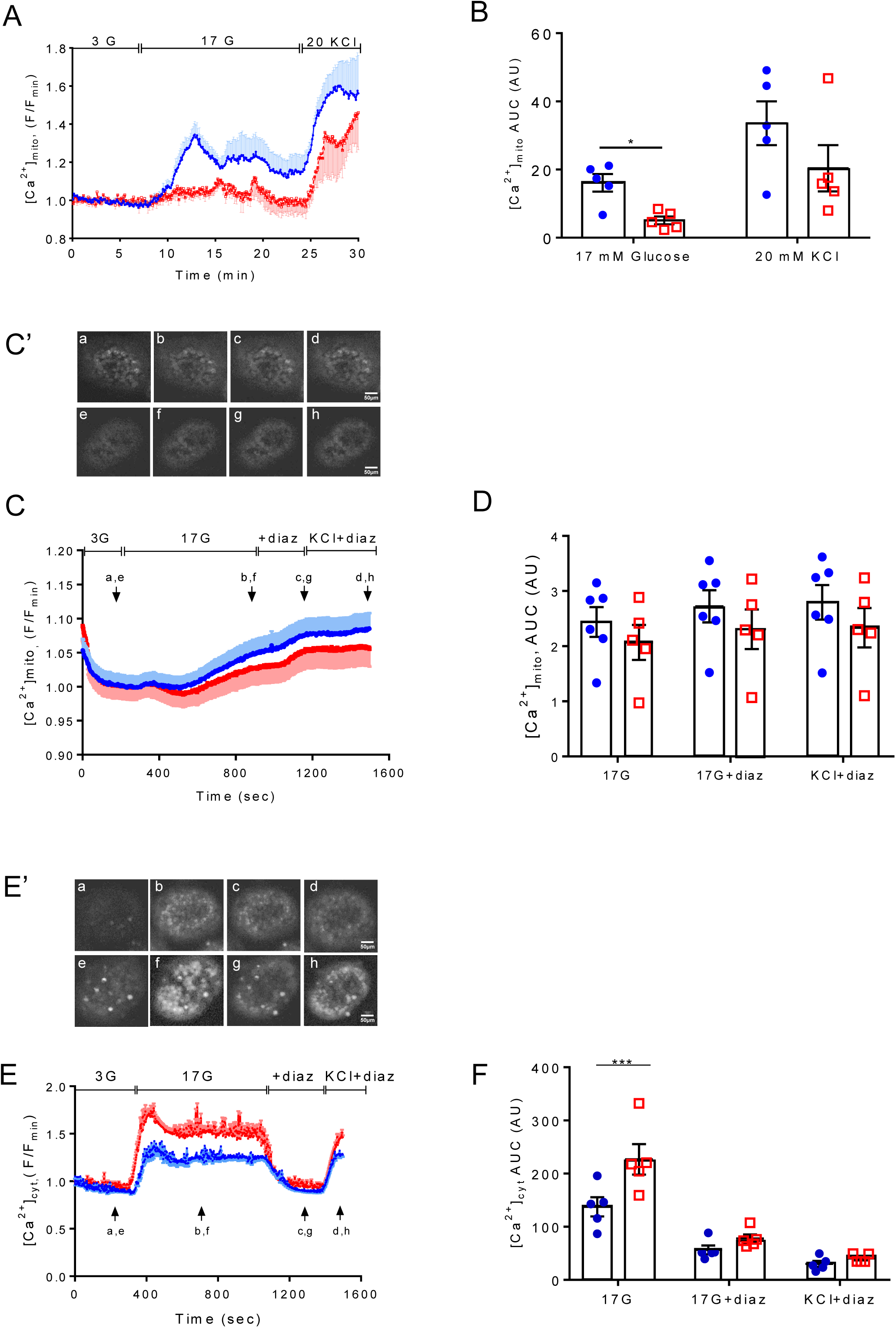
MCU deletion from pancreatic β-cells diminishes *in vitro* mitochondrial Ca^2+^ uptake in dissociated islets but not cytoplasmic [Ca^2+^] in whole islets. (A) Genetically-encoded recombinant Ca^2+^ probe, R-GECO, was used to assess mitochondrial Ca^2+^ dynamics in response to 17 mM glucose and 20 mM KCl in dissociated β-cells. (B) AUC corresponding to the data shown in (A): *p<0.05; *n=*5 trials, 3 mice per genotype; data points correspond to individual trials; 8-15 min for stimulation with 17 mM glucose. (C’) Each snapshot of isolated WT (upper panel) and KO (lower panel) islets was taken during the time points shown with an arrow. See also supplemental movies Pericam WT and Pericam KO. (C) Mitochondrial Ca^2+^ changes in response to 17 mM glucose (with or without diazoxide) and 20 mM KCl were assessed following mito Pericam infection. Traces represent mean normalised fluorescence intensity (F/F_min_) over time. Scale bar= 50μm. (D) AUC corresponding to the data shown in (C): *n=*5-6 trials, 3 mice per genotype; data points correspond to individual trials; no significant differences detected. (E’) Each snapshot of isolated WT (upper panel) and KO (lower panel) islets was taken during the time points shown with an arrow. See also supplemental movies Cal-520 WT and Cal-520 KO. (E) Cytoplasmic Ca^2+^ changes in response to 17 mM glucose (with or without diazoxide) and 20 mM KCl were assessed following Cal-520 uptake. Traces represent mean normalised fluorescence intensity (F/F_min_) over time. Scale bar= 50μm. (F) AUC corresponding to the data shown in (E): ***p<0.001; *n=*5 trials, 3 mice per genotype; data points correspond to individual trials. Islets were isolated from 8-10 week old male mice maintained on standard chow diet. Values represent mean ± SEM. AU, arbitrary unit; AUC, area under the curve. Statistical analyses were performed using two-way ANOVA tests and Sidak’s correction for multiple comparisons.

*Mcu* deletion attenuated glucose-stimulated increases in [Ca^2+^]_mt_ in dissociated islets (Fig. 2A-B), as demonstrated within normalised traces and by determination of the mean area under the curve at 17mM glucose (AUC; p<0.05). The [Ca^2+^]_mt_ increase in response to depolarisation with KCl was largely maintained in βMCU-KO. The [Ca^2+^]_mt_ was also assessed in whole islets using mito Pericam where a trend towards reduced mitochondrial Ca^2+^ uptake was observed (Fig. 2C-D). To determine whether the impaired [Ca^2+^]_mt_ changes in βMCU-KO β-cells may simply reflect altered cytosolic Ca^2+^ ([Ca^2+^]_cyt_) dynamics, the latter were also explored using the Ca^2+^-sensitive dye (Cal-520) in whole islets. Interestingly, [Ca^2+^]_cyt_ increases in whole βMCU-KO islets were significantly increased in response to glucose in comparison to WT animals (AUC; p<0.001; Fig. 2E, F).

### MCU deletion from pancreatic β-cells reduces mitochondrial ATP production and cytosolic accumulation whereas mitochondrial membrane depolarisation decreases in response to glucose

Given the significant reduction in GSIS observed in βMCU-KO islets *in vitro* and despite improved cytosolic Ca^2+^ dynamics, we next sought to determine whether an alteration in glucose metabolism might contribute to the attenuated insulin secretion. To address this, we utilised real-time fluorescence imaging of the ATP sensor, Perceval (9). A rise in the ATP:ADP ratio was prompted in control β-cells by a step increased in glucose from 3 to 17 mM (9). This change was significantly blunted in *Mcu* null β-cells (AUC; p<0.05; Fig. 3A, B). This was accompanied by a potentiation in the increase in mitochondrial membrane potential (Δψ_m_), as assessed by monitoring TMRE fluorescence in βMCU-KO mouse β-cells (AUC_700-720s_; p<0.05; Fig. 3C, D).

**Figure 3.**
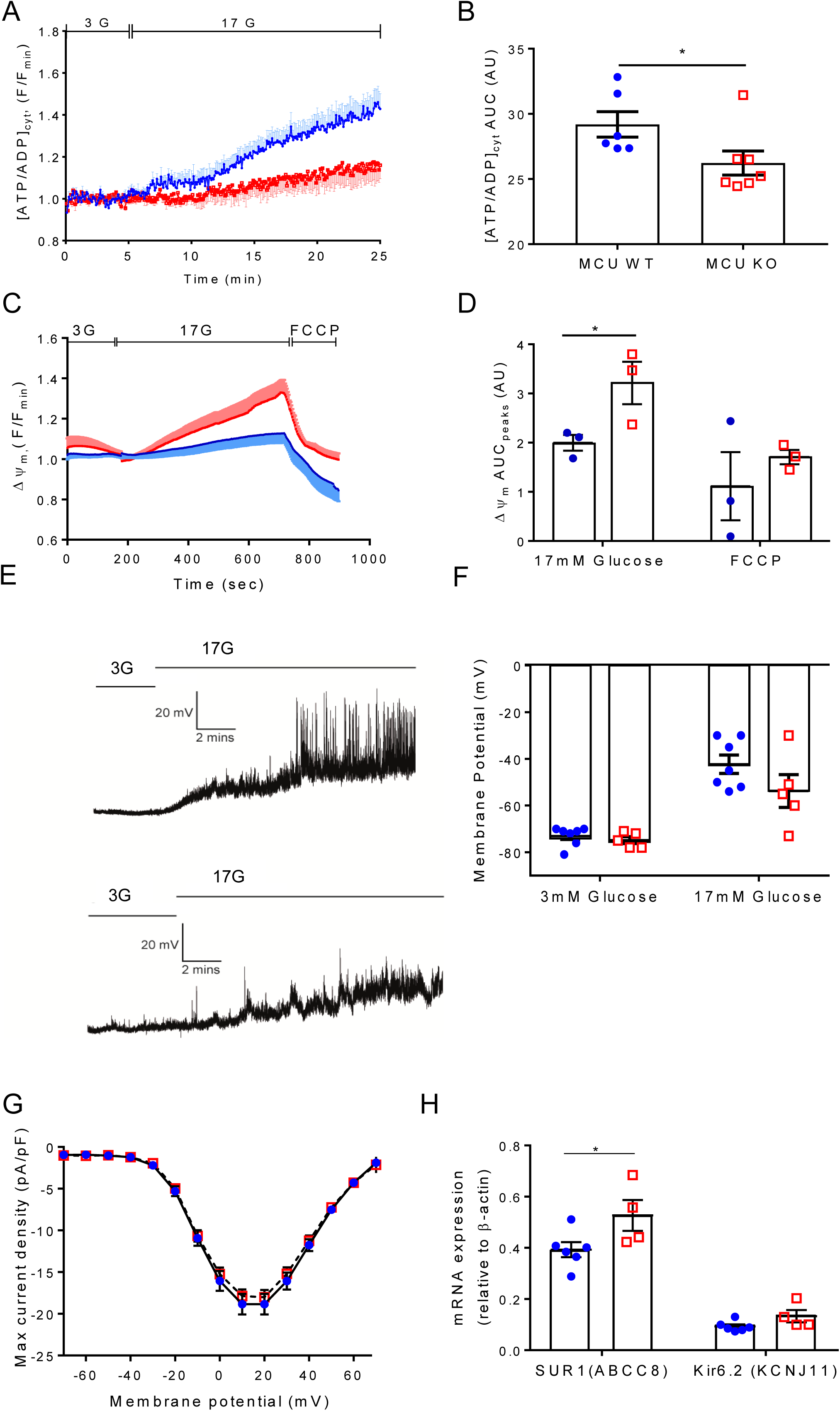
MCU ablation from pancreatic β-cells diminishes ATP production and mitochondrial membrane depolarisation in response to high glucose. (A) Changes in the ATP:ADP ratio in response to 17 mM glucose was examined in dissociated β-cells using the ATP sensor Perceval. (B) AUC values corresponding to (A) (p<0.05, *n=*6-7 trials, 3 mice per genotype; unpaired two-tailed Student’s t-test). (C) Cells were loaded with TMRE to measure changes in Δψ_m_, and perifused with 3 mM, 17 mM glucose or 1μM FCCP as indicated. Traces represent normalised fluorescence intensity (F/F_min_) over time. (D) AUC were determined from the data shown in (C): AUC_700-720s_ peak at 17mM glucose (p<0.05) and presented as mean of the values ± SEM. Data points are from *n=*3 mice per genotype (two trials per mouse). (E) Representative current-clamp recordings of individual β-cells from WT (upper trace) and βMCU-KO mice (lower trace) displaying the membrane potential response from 3 to 17 mM glucose. (F) Mean membrane potential responses (*n=*5-7 trials, 3 mice per genotype; two-way ANOVA test and Sidak’s multiple comparisons test). (G) Activation of β-cell VDCCs in response to 17mM glucose and indicated voltage steps (*n=*23-24 islets, 3 mice per genotype). (H) qRT-PCR quantification of *Kcnj11* and *Abb8* expression (*p<0.05; *n=*4-6 mice per genotype; unpaired two-tailed Student’s t-test and Mann Whitney correction). Islets were isolated from 8-10 week old male mice maintained on standard chow diet. Values represent mean ± SEM. AU, arbitrary unit; AUC, area under the curve.

Altered ATP production in response to high glucose is expected to affect the activity of K_ATP_ channels (27). Assessed in single β-cells using perforated patch-clamp electrophysiology (9), the extent of membrane potential depolarisation in response to a step increase in extracellular glucose from 3 to 17 mM did not differ significantly between βMCU-KO and control β-cells, although there was a trend towards weaker depolarisation in KO cells (Fig. 3E, F). VDCC currents, measured by whole-cell voltage clamp (22), displayed no apparent differences in response to 17 mM glucose (Fig. 3G). Interestingly expression of the K_ATP_ channel subunit *Kcnj11* was significantly elevated in βMCU-KO islets (AUC; p<0.05; Fig. 3H) any may contribute to reduced electrical activity (Fig. 3E, F).

### Lowered β-cell mass in MCU-deleted mice

Analysis using optical projection tomography (OPT; Fig. 4A) revealed that pancreata from βMCU-KO mice displayed decreased numbers of islets at the extremes of the size spectrum (p<0.01) in comparison to WT mice (Fig. 4B) and a decrease in total β-cell volume (AUC; p<0.01; Fig. 4C).

**Figure 4.**
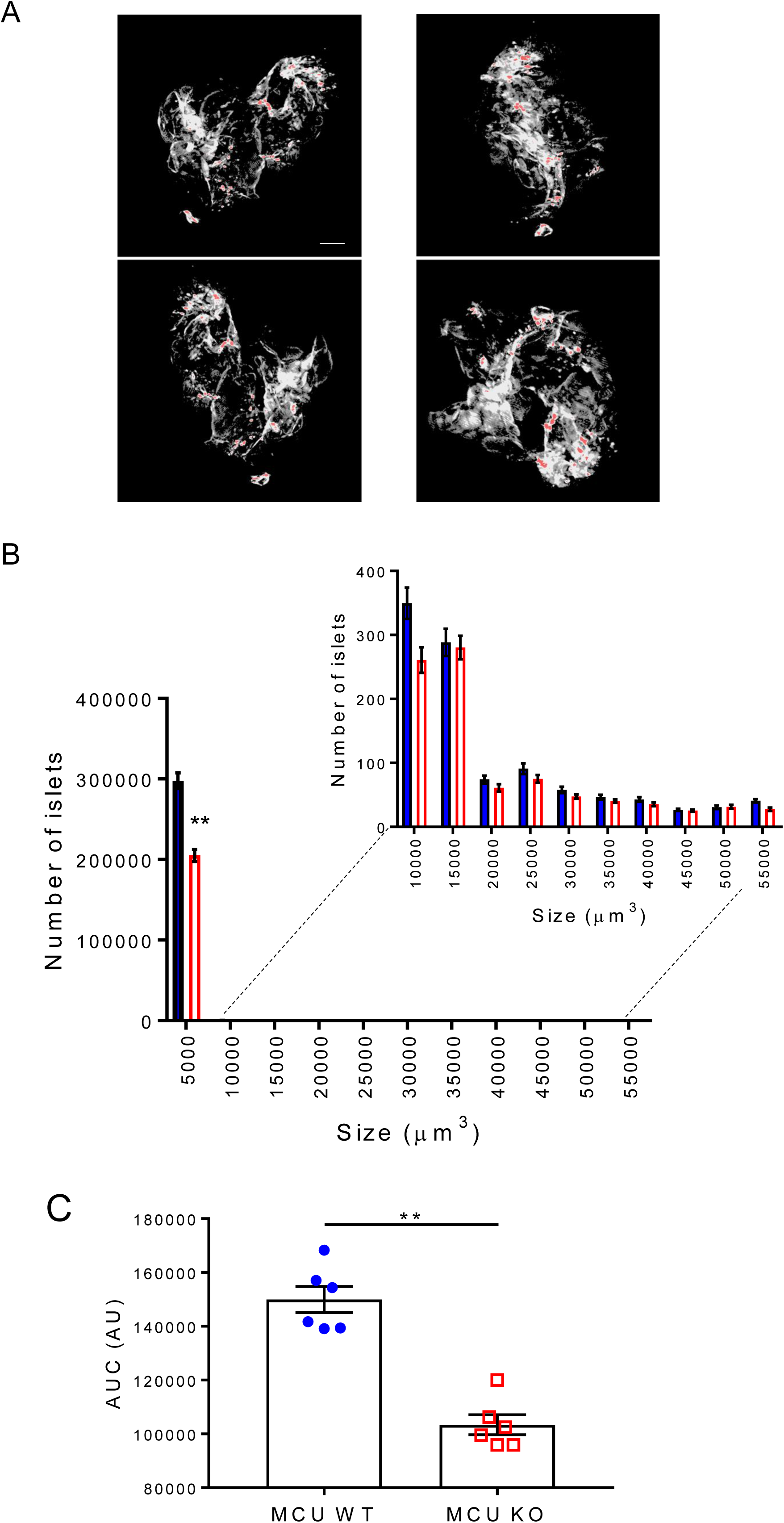
Effect of *Mcu* deletion on β-cell mass. (A) Optical projection tomography shows images of representative pancreata stained with insulin (pseudo-colour, red) to indicate islets of different sizes. Scale bar=500μm. (B) Quantification of the number of islets indicates a significant (p<0.01) decrease in smaller islets in βMCU-KO mice (*n=*6 animals per genotype). (C) Changes in overall β-cell mass (Area under the curve, AUC; p<0.01; unpaired two-tailed Student’s t-test and Mann Whitney correction, *n=*6 animals per genotype). Values represent mean ± SEM.

### Loss of MCU from pancreatic β-cells does not alter body mass or fed glycaemia but impairs GSIS *in vivo*

We next explored the role of β-cell MCU in the control of insulin secretion and *in vivo* glucose homeostasis. βMCU-KO animals displayed normal growth and weight changes from 6 to 24 weeks (**Fig. S1A**). However, male βMCU-KO mice showed a slight, but significant, increase (p<0.05) in weight gain from 20-24 weeks. We observed no differences in random fed blood glucose levels between βMCU-KO and control animals at all ages examined (**Fig. S1B**). No genotype-dependent differences in the above metabolic parameters were observed in female mice at any age (**Fig. S1C, S1D**; **Fig. S2A-D**).

Glucose tolerance was investigated in βMCU-KO and WT mice by intraperitoneal injection of 1g/kg body weight glucose (IPGTT) at 8, 12, 16 and 24 weeks of age. A small but significant improvement in glucose tolerance was observed in male βMCU-KO mice *vs* controls at 8 (p<0.001) or 12 (p<0.05) weeks of age (Fig. 5A, B). Older male animals displayed unaltered glucose tolerance (Fig. 5C, D) *vs* WT mice.

**Figure 5.**
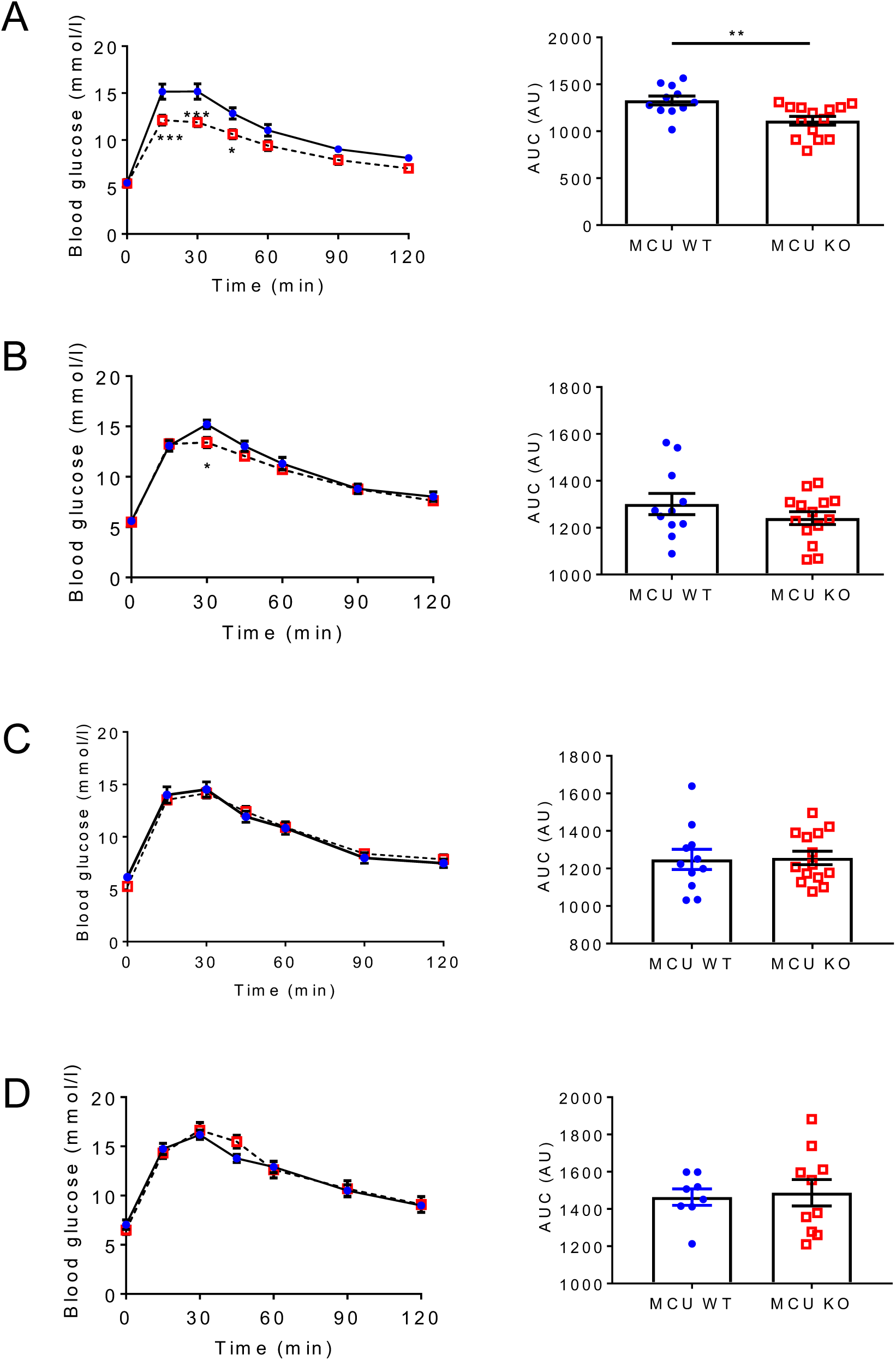
Male βMCU-KO mice display slightly improved glucose tolerance. Glucose tolerance was measured in male MCU-KO and littermate control (WT) mice by intraperitoneal injection of glucose (1g/kg body weight) at (A) 8, (B) 12, (C) 16 and (D) 24 weeks of age. The AUC is shown to the right of each graph (p<0.05 or p<0.001, *n=*8-14 mice per genotype).

To assess GSIS *in vivo*, 8-10 week-old male mice were challenged with 3g/kg glucose and plasma insulin sampled at 0, 5, 15 and 30 min. (Fig. 6A, B). Although improved glucose tolerance was observed in βMCU-KO animals post 15 min. IP administration (Fig. 6A), a dramatic reduction (p<0.001) in insulin release was observed 5 min. post-glucose injection (Fig. 6B). βMCU-KO animals also displayed improved oral glucose tolerance at 15 and 30 min. post-oral gavage (Fig. 6C; **p<0.05),** increased insulin secretion at 5 min. (Fig. 6D; **p<0.05)** vs WT littermates. No differences in intraperitoneal insulin tolerance (Fig. 6 E, F) or c-peptide levels (not shown) were observed between WT and KO mice.

**Figure 6.**
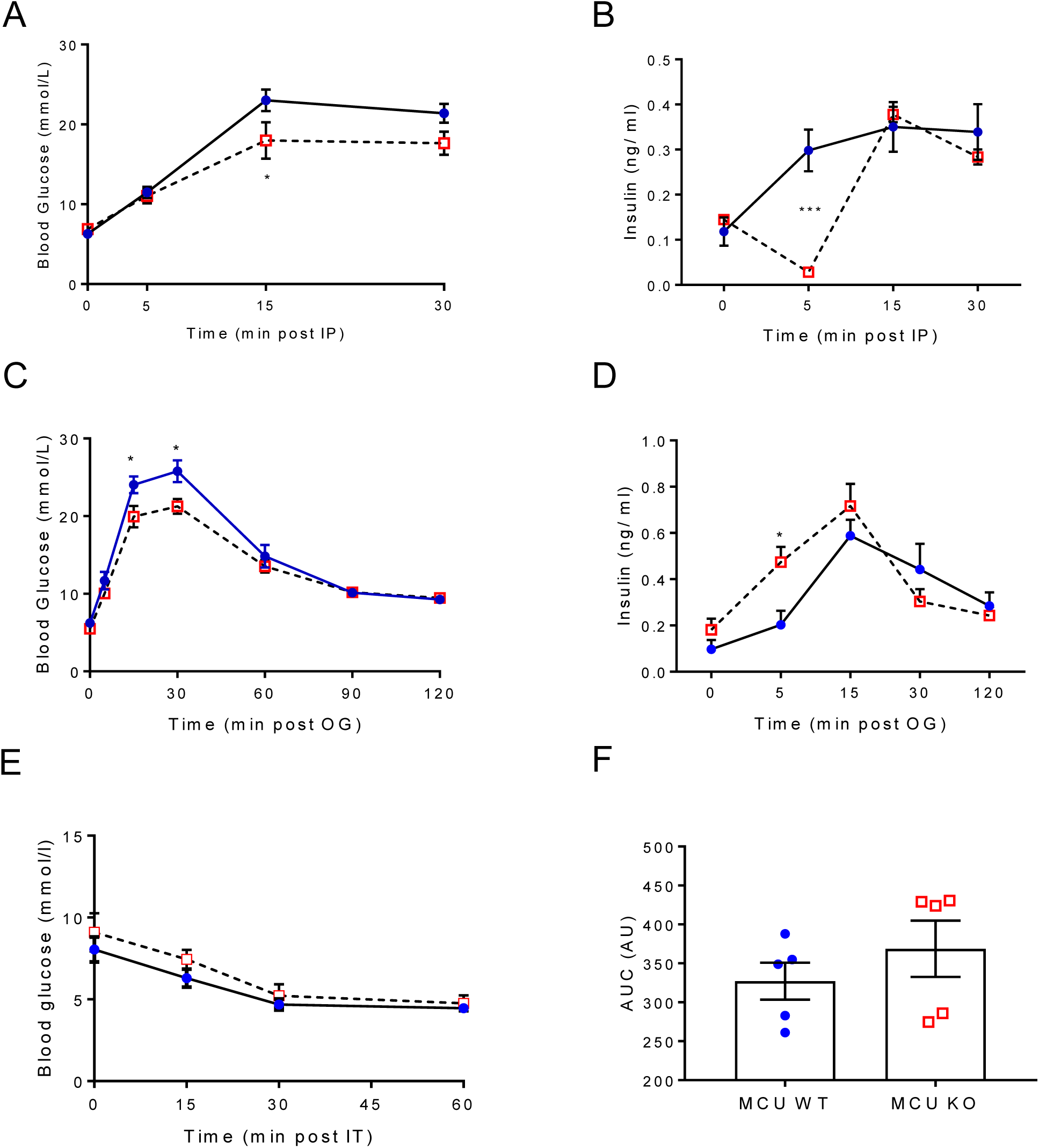
βMCU-KO mice display enhanced glucose tolerance following intraperitoneal or oral gavage glucose administration. (A) Glycaemia and (B) glucose (3 g/kg body weight)-induced insulin secretion were assessed in βMCU-KO and WT mice (p<0.05 or p<0.001; 8-10 weeks old, *n=*6-9 mice per genotype). (C) Plasma glucose and (D) insulin, during the oral glucose tolerance test in βMCU-KO and WT mice (p<0.05; *n=*7-9 per genotype). (E) Challenging 8-10 week-old male mice with a 0.75 U/kg body weight insulin injection showed normal insulin sensitivity. (F) The AUC is also shown (*n=*5 mice per genotype). All mice were maintained on a standard chow diet. Values represent mean ± SEM. AU, arbitrary unit; AUC, area under the curve.

Finally, to impose a metabolic stress, βMCU-KO mice and control littermates were maintained on a HFHS diet and subjected to the same tests as in (Fig. 6A, B). Again, no differences in glycaemia or insulin secretion were observed between phenotypes (**Fig. S3A, B**). No genotype-dependent differences in c-peptide secretion were apparent (not shown).

## Discussion

Mitochondria are highly dynamic organelles which play an important role in maintaining normal β-cell function and secretory responses to glucose (28–30). As mitochondrial dynamics and biogenesis are impaired in the face of insulin resistance and in type 2 diabetes in these cells (8), it is conceivable that preserving the normal function of these organelles may slow the loss of normal insulin secretion and disease progression (31; 32).

Recent findings (10) have demonstrated the importance of MCU for Ca^2+^ uptake into mitochondria in several mammalian cell types, and established this as the most important route for Ca^2+^ accumulation into these organelles. In the present studies, efficient deletion of both MCU isoforms was achieved by targeting exons 11 and 12 of the *Mcu* gene using recombination with efficient and selective *Ins1*Cre deleter strain. This resulted in near complete elimination of a functional *Mcu* gene throughout the β-cell population *in vivo*. In agreement with previous studies using RNA silencing (9; 15), *Mcu* deletion attenuated GSIS and mitochondrial Ca^2+^ uptake in response to high glucose in dissociated islets (9). The response to depolarisation by KCl was less markedly affected, possibly reflecting the opening of other mitochondrial Ca^2+^ transporters/channels at high cytosolic [Ca^2+^] levels (33). Alternative mitochondrial Ca^2+^ entry pathways may involve ryanodine receptors, as observed on the inner mitochondrial membrane in neurons (34), or the rapid mode of mitochondrial Ca^2+^ uptake (RAM) in the liver. The existence of these pathways has not, as yet, been demonstrated in pancreatic β-cells.

Surprisingly, quantification of [Ca^2+^]_cyt_ in intact islets demonstrated larger increases in response to high glucose in KO islets (Fig. 7A,B), perhaps reflecting an impact on Ca^2+^ oscillation frequency, β cell-β cell communication and three-dimensional electrical communication through gap junctions (35). The sharp decrease in GSIS *in vitro* in the face of higher cytosolic [Ca^2+^] (Fig. 7B) is, again, paradoxical and argues that lowered ATP:ADP, alongside impairments in amplifying processes (36), such as the Ca^2+^-dependent intra-mitochondrial generation of putative coupling molecules such as glutamate (37), or others (2), exert a dominant, inhibitory effect on secretion in KO mice (see below).

**Figure 7.**
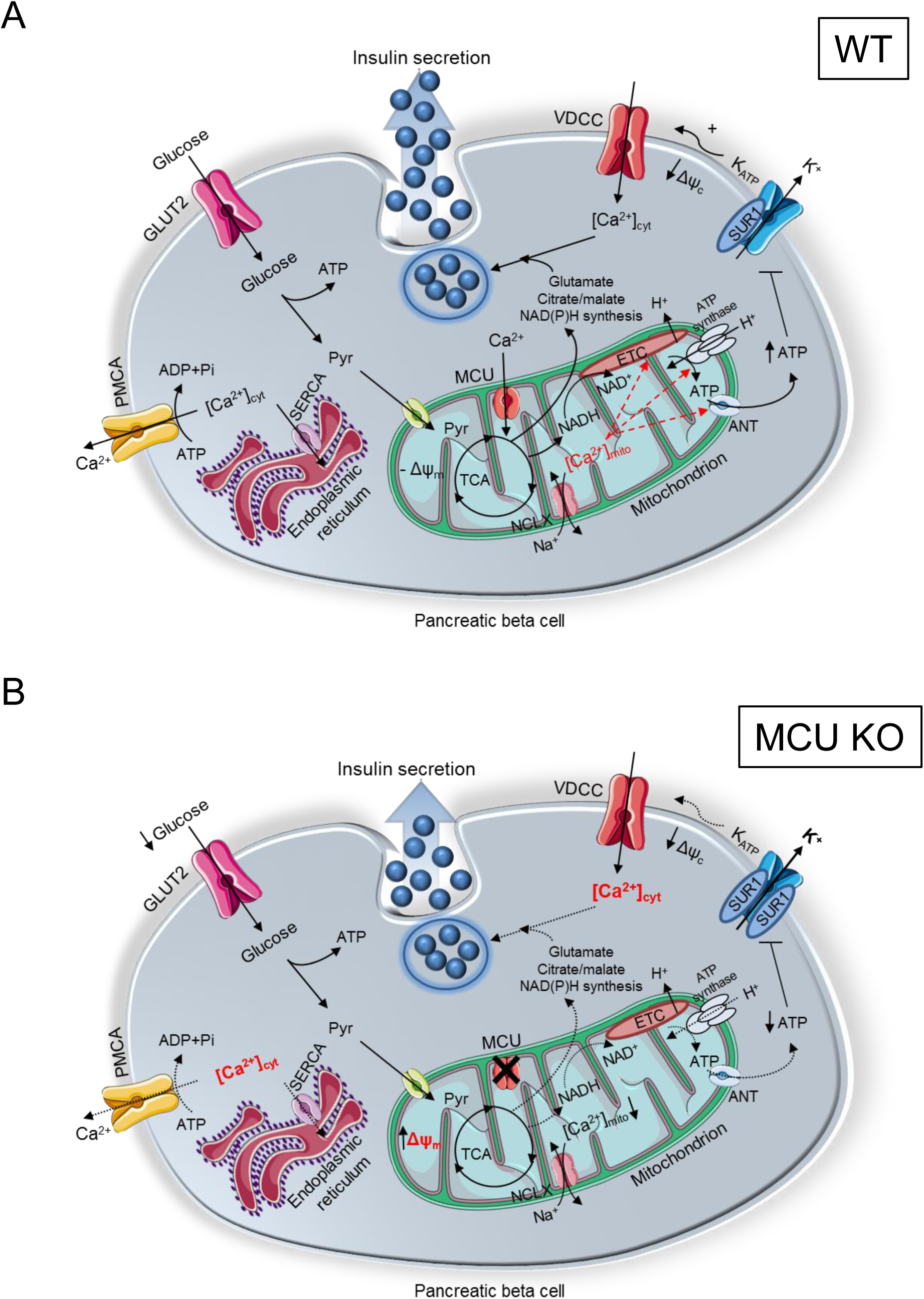
Putative involvement of ΜCU in coordinating the response of β-cells to nutrient supply, and impact of MCU deletion on GSIS. (A) Glucose is taken up by β-cells and catabolised glycolytically. The formed pyruvate is metabolised by mitochondria through the citrate (TCA) cycle, leading to an increased mitochondrial proton motive force (hyperpolarised Δψ_m_) and accelerated ATP synthesis. By entering mitochondria via the MCU, Ca^2+^ potentiates oxidative metabolism to counter-balance ATP consumption. Ca^2+^ exits mitochondria via NCLX. Consequently, the cytoplasmic ATP:ADP ratio rises, which causes further closure of ATP-sensitive K^+^ channels, depolarisation of the plasma membrane potential (Δψ_c_), opening of VDCCs, and influx of Ca^2+^. Elevated [Ca^2+^]_cyt_ triggers a number of ATP-dependent processes including insulin secretion and Ca^2+^ removal into the ER (sarco(endo)plasmic reticulum Ca^2+^ ATPase; SERCA) and extracellular medium (plasma membrane Ca^2+^ ATPase, PMCA). Mitochondrial metabolism is also activated by amino acids such as glutamate, citrate/malate which appear to be necessary for appropriate generation of regulatory, “amplifying” signals for insulin secretion. (B) Following MCU deletion, [Ca^2+^]_mito_ is reduced leading to a more highly polarised Δψ_m_, weaker oxidative or amino acid metabolism and ATP synthesis, perhaps due to a decrease in mitochondrial F_1_F_0_ATPase and/or adenine nucleotide transferase (ANT) activity. This is expected to result in result less closure of ATP-sensitive K^+^ channels (K_ATP_, further potentiated by increased expression of the SUR1 subunit, weaker plasma membrane depolarisation and Ca^2+^ influx. Importantly, lowered ATP supply to the cytosol is expected to restrict Ca^2+^ pumping across the plasma membrane, as well as into the ER. Despite reporting elevated [Ca^2+^]_cyt_ in βMCU-KO mice, insulin secretion *in vitro* was impaired possibly due to lower Ca^2+^ dependent intra-mitochondrial generation of putative coupling molecules such as glutamate, citrate/malate.

Importantly, our observations support the view that Ca^2+^ accumulation by mitochondria stimulates ATP consumption, consistent with a reduction in glucose-stimulated ATP:ADP ratio in the KO mouse (6). Additionally, mitochondrial membrane potential was increased, particularly in glucose-stimulated conditions, in dissociated islets from KO *vs* WT mice. An increase in Δψ_m_ in the face of lowered cytosolic ATP:ADP is consistent with a decrease in F_1_/F_0_ATPase activity (38) (Fig. 7B), an enzyme previously reported to be Ca^2+^ regulated in other tissues (8). Reduced mitochondrial Ca^2+^ extrusion via flux through the mitochondrial Na^+^/Ca^2+^ exchanger NCLX (electrogenic: 3Na^+^: 1Ca^2+^) may also contribute (39). Additionally, lowered cytosolic ATP is expected to restrict Ca^2+^ pumping across the plasma membrane (Ca^2+^ATPase), as well as into the endoplasmic reticulum (SERCA; Fig. 7B). Of note, despite attenuated ATP increases and greater accumulation of cytosolic Ca^2+^, we observed only a trend towards lower glucose-stimulated plasma membrane depolarisation and no significant difference in VDCC activity between WT and KO β-cells. Quantification of K_ATP_ channel subunit expression revealed elevated *Kcnj11* mRNA levels in βMCU-KO mice which, alongside lowered cytosolic ATP:ADP increases, is expected to lower membrane excitability (Fig. 7B) and Ca^2+^ entry (40).

In addition to the above functional alterations, βMCU-KO mice displayed decreased β-cell mass. This may reflect either impaired proliferation or generation from progenitor cells in the absence of functional *Mcu*, or altered cell death (20). Of note, Zhao et al., (41) have recently reported that down-regulation of MCU enhances autophagic death in neurons due to the activation of AMP-activated protein kinase (AMPK), a known regulator of β-cell mass (42).

Extending to the *in vivo* setting, the current and earlier (9; 15) *in vitro* data demonstrating roles for MCU in the control of glucose-induced insulin secretion, we show that insulin secretion is impaired in βMCU-KO *vs* control mice 5 min. post to intraperitoneal injection of glucose. Surprisingly, however, insulin secretion post-oral gavage was elevated in KO mice at 5 min. perhaps suggesting that gut-derived factors such as the gastric inhibitory polypeptide and glucagon-like peptide-1 (GIP and GLP-1) are partially responsible for the enhanced glucose-stimulated insulin secretion from the β-cells (43). Future studies will be necessary to explore this possibility.

Interestingly, earlier studies inactivating MCU globally in the mouse, or in selected tissues, have consistently reported relatively minor phenotypes (44). Global MCU null mice displayed relatively unimpaired cardiac and skeletal muscle function and respiration (11) despite a near-complete ablation of Ca^2+^ accumulation by mitochondria in these cells. The present results are thus in line with these earlier findings. It is still unknown why insulin secretion is acutely impaired 5 min. post intraperitoneal glucose administration in KO mice but compensatory mechanisms are likely to emerge after that period to maintain normal cellular energy homeostasis and signalling *in vivo*. One intriguing possibility, which may be of particular relevance to the nutrient-responsive β-cell, is that the mitochondrial Na^+^-Ca^2+^ exchanger NCLX operates in the reverse mode at low Δψ_m_, thus allowing Ca^2+^ influx (45) to provide a compensatory mechanism for the loss of MCU.

The findings here also provide evidence of a role for additional, mitochondrially-derived metabolic signals whose generation depends upon mitochondrial Ca^2+^ uptake and which serve to potentiate the actions of increased cytosolic Ca^2+^. Such molecules have been proposed to underlie the “amplification” (K_ATP_-channel independent) component of glucose-stimulated insulin secretion (36) but still remain elusive. Recent studies have focused on mitochondrial pathways of glucose metabolism, and the generation of second messengers other than ATP that such pathways might generate. Islet β-cells express both pyruvate carboxylase (PC) and pyruvate dehydrogenase (PDH) in abundance, such that, in the fed state, pyruvate flows into mitochondrial metabolic pathways in roughly equal proportions through the anaplerotic and oxidative entry points (2; 46). Impairments in pyruvate/isocitrate cycle activity, and the generation of its by-products (notably α-ketoglutarate (α-KG) and NADPH) might therefore restrict normal insulin secretion in βMCU-KO mice (Fig. 7B).

## Conclusion

To the best of our knowledge, this study provides only the second description of a conditional null *Mcu* KO mouse and reveals a critical role for MCU-mediated mitochondrial Ca^2+^ influx in the pancreatic β-cell *in vitro* and *in vivo*. The mechanisms which compensate *in vivo* for defective insulin secretion in βMCU-KO mice, ensuring near-normal or improved glucose homeostasis will need further exploration in the future.

Our findings suggest that changes in MCU expression or activity may contribute to defective insulin secretion in some forms of diabetes. An alteration in the ratio of the active (MCUa) form of the channel versus MCUb, a dominant-negative form of the carrier (26), might also play a part in the disease process in some settings.

## Supporting information

On line appendix

## Conflict of interest

The authors declare that they have no conflicts of interest with the contents of this article.

## Funding

G.A.R. was supported by a Wellcome Trust Senior Investigator Award (WT098424AIA) and Investigator Award (212625/Z/18/Z), MRC Programme grants (MR/R022259/1, MR/J0003042/1, MR/L020149/1) and Experimental Challenge Grant (DIVA, MR/L02036X/1), MRC (MR/N00275X/1), Diabetes UK (BDA/11/0004210, BDA/15/0005275, BDA16/0005485) and Imperial Confidence in Concept (ICiC) grants, and a Royal Society Wolfson Research Merit Award. I.L. was supported by Diabetes UK Project Grant 16/0005485. This project has received funding from the Innovative Medicines Initiative 2 Joint Undertaking under grant agreement No 115881 (RHAPSODY). This Joint Undertaking receives support from the European Union’s Horizon 2020 research and innovation programme and EFPIA. This work is supported by the Swiss State Secretariat for Education, Research and Innovation (SERI) under contract number 16.0097. RR was supported by grants from the Italian Ministries of Health (Ricerca Finalizzata.) and of Education, University and Research (FIRB), the European Union (ERC mitoCalcium, no. 294777), the National Institutes of Health (grant #1P01AG025532-01A1), the Italian Association for Cancer Research (AIRC IG18633), and Telethon-Italy (GGP16029).

## Author contributions

E.H, E.G. and G.dSX performed experiments and analysed data. F.S. and M.C.C. generated and amplified Perceval and R-GECO viruses. I.L. and A.M.S. were responsible for the maintenance and genotyping of mouse colonies. T.J.P. examined RNAseq data and designed experiments. R.R. provided reagents. G.A.R. designed the study and wrote the manuscript with E.H. and E.G., with input from all authors. G.A.R. is the guarantor of this work and, as such, had full access to all the data in the study and takes responsibility for the integrity of the data and the accuracy of the data analysis.

## Data and Resource Availability

All data generated or analyzed during this study are included in the published article (and its online supplementary files). No applicable resources were generated or analyzed during the current study.

## Prior presentation

Some of this work has been presented at the following conferences: as a poster at the EASD Annual conference, Lisbon, Portugal, September 2017, with an accompanying abstract in Haythorne, E.A., Martinez-Sanchez, A., Rizzuto, R., and Rutter, G.A. The mitochondrial uniporter (MCUa) is required for glucose-stimulated mitochondrial Ca^2+^ uptake and insulin secretion in mouse pancreatic beta cells, *Diabetologia*, **60**, Supp1, S1-S608. 431 (2017): as a poster at the Diabetes UK Annual Professional Conference, London, U.K., March 2018, with the accompanying abstract: Rutter, G.A. Haythorne, E.A., Georgiadou, E., Da Silva Xavier, G., Pullen, T.J., Rizzuto, R., Martinez-Sanchez, A., McGinty, J.A. and French, P.M. Pancreatic beta cell-selective deletion of the mitochondral Ca^2+^ uniporter MCU impairs glucose stimulated insulin secretion in vitro but not in vivo. *Diabetic Med* 35:42 (2018). Data were also included in an oral presentation at a conference entitled “Mitochondrial Form and Function” at University College London, U.K., September 2017.

## Abbreviations

[Ca^2+^]_cyt_: Cytoplasmic Ca^2+^ concentration
[Ca^2+^]_mt_: Mitochondrial Ca^2+^ concentration
ADP: Adenosine diphosphate
AMPK: Adenosine monphosphate-activated protein kinase
ANT: adenine nucleotide transferase
ATP: Adenosine triphosphate
AUC: Area under curve
FCCP: Carbonyl cyanide-4-phenylhydrazone
GIP: Gastric inhibitory polypeptide
GLP-1: Glucagon-like peptide-1
GSIS: Glucose-stimulated insulin secretion
HTRF: homogeneous time-resolved fluorescence-based assay
IPGTT: Intraperitoneal glucose tolerance test
KATP: ATP-sensitive K^+^ (K_ATP_) channels
KCl: Potassium chloride
KO: Knockout
MCU: Mitochondrial Ca^2+^ uniporter
NCLX: Na^+^-Ca^2+^ exchanger
OGTT: Oral gavage tolerance test
PC: Pyruvate carboxylase
PDH: Pyruvate dehydrogenase
RAM: Rapid mode of mitochondrial Ca^2+^ uptake
SERCA: Sarco/endoplasmic reticulum Ca^2+^-ATPase
T2D: Type 2 diabetes
TCA: tricarboxylic acid
TMRE: Tetramethylrhodamine ethyl ester
VDCCS: Voltage-dependent Ca^2+^ channels
WT: Wild type
βMCU-KO: Mitochondrial Ca^2+^ uniporter null animal in the beta cell
α-KG: α-ketoglutarate
Δψ_m_: Mitochondrial membrane potential

