## Supplementary material for "The mitochondrial Ca^2+^ uniporter MCU is required for normal glucose-stimulated insulin secretion *in vitro* and *in vivo*": On line appendix

**Supplemental Data and Methods**

**Supplemental Table 1.** Sequences of primers used for genotyping of MCUa Flox. (F: forward, R: reverse).

| | |
|---|---|
| *MCUa* | F: 5’CTGCTTCTGTGTACATTCAAGGATG |
| | R: 5’CTCGGTTCTAGATACTGGCATTCAC |

**Supplemental Table 2.** List of primers used for qRT-PCR amplification (F: forward, R: reverse).

| | | |
|---|---|---|
| qRT-PCR | *MCUa* | F-GAGCAGCATCAGCTTAACAAAGAG |
| | | R-TCTGCTAATTTCAATTCGTACCTTCTC |
| | *β-actin* | F-CGAGTCGCGTCCACCC |
| | | R-CATCCATGGCGAACTGGTG |
| | *ABCC8* | *F-GCCTACGCATCTCAGAAACCA* |
| | | *R-CCATCTTGTACCTTTGCTTATTGAAG* |
| | *KCNJ11* | *F-CACGGCGGGATAAGTCTACCT* |
| | | *R-AATCATTTGCCCCCTTCTTGT* |

**Supplemental Figures and Legends**

**Supplemental Figure 1. Male but not female βMCU-KO mice display slightly increased weight gain on standard chow diet, compared to littermate controls.** (A) Body weight and (B) random fed glycaemic profile of male and (C, D) female βMCU-KO and littermate control mice (WT). (A, B) $p<0.05$ by two-way ANOVA and Bonferroni correction for multiple tests, n=11-14 mice per genotype) and (C, D) n=10-12 mice per genotype) accordingly. Blue circles, WT and red squares, βMCU-KO mice.

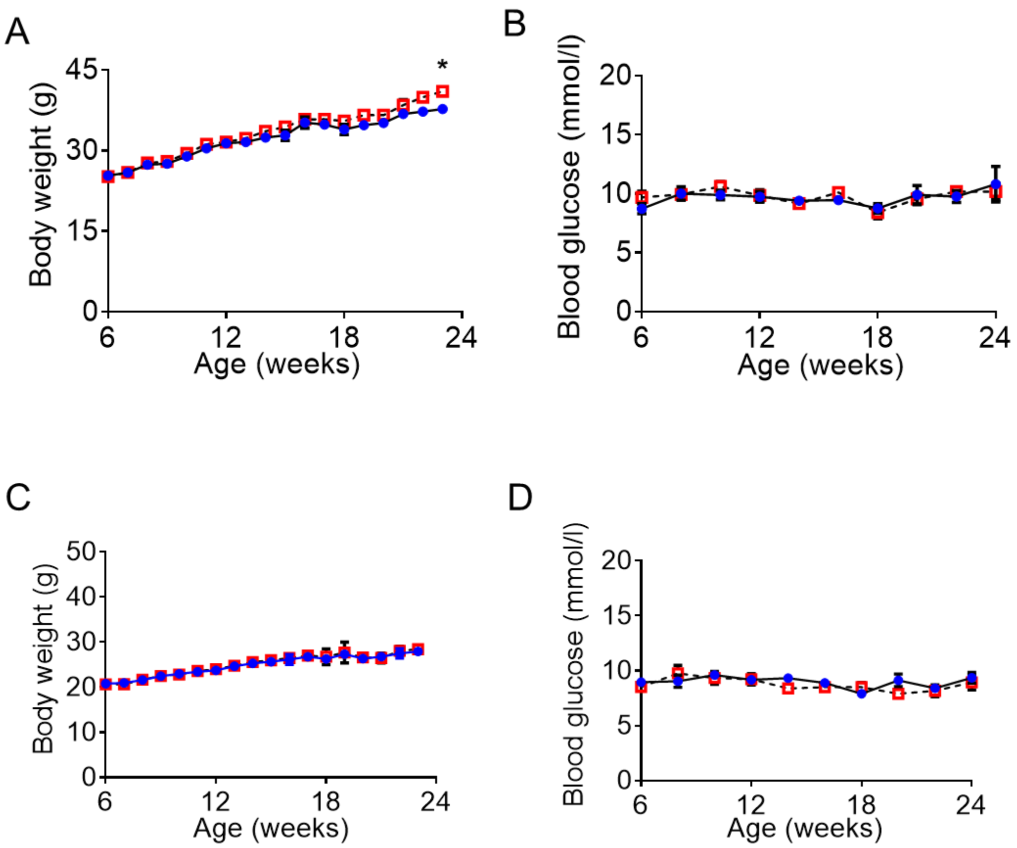

**Supplemental Figure 2. Female βMCU-KO mice display normal glucose tolerance.** (A) Glucose tolerance was measured in female βMCU-KO and littermate control (WT) mice by intraperitoneal injection of glucose (1g/kg body weight) at 8, (B) 12, (C) 16 and (D) 24 weeks of age. Statistical analysis by two-way ANOVA and Bonferroni correction for multiple tests. The AUC is shown to the right of each graph (n=10-12 mice per genotype). All mice were maintained on a standard chow diet. Values represent mean ± SEM. AU, arbitrary unit; AUC, area under the curve. Statistical significance determined by unpaired Student's t-test.

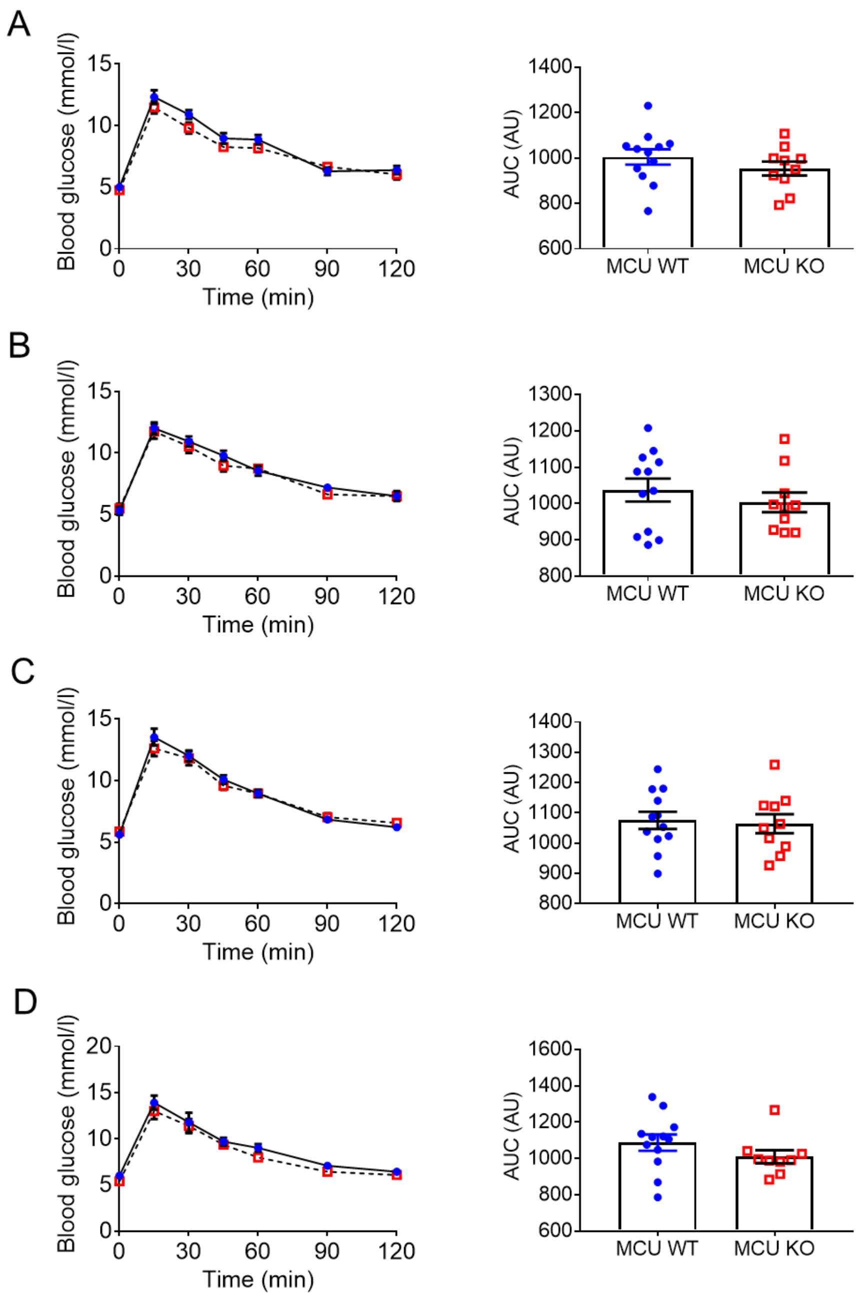

**Supplemental Figure 3. Male βMCU-KO mice display normal glucose tolerance and insulin secretion following a HFHS diet.** (A) Glycaemia and (B) glucose (3 g/kg body weight)-induced insulin secretion were assessed in βMCU-KO and WT mice (8 weeks old, n=5-7 mice per genotype).

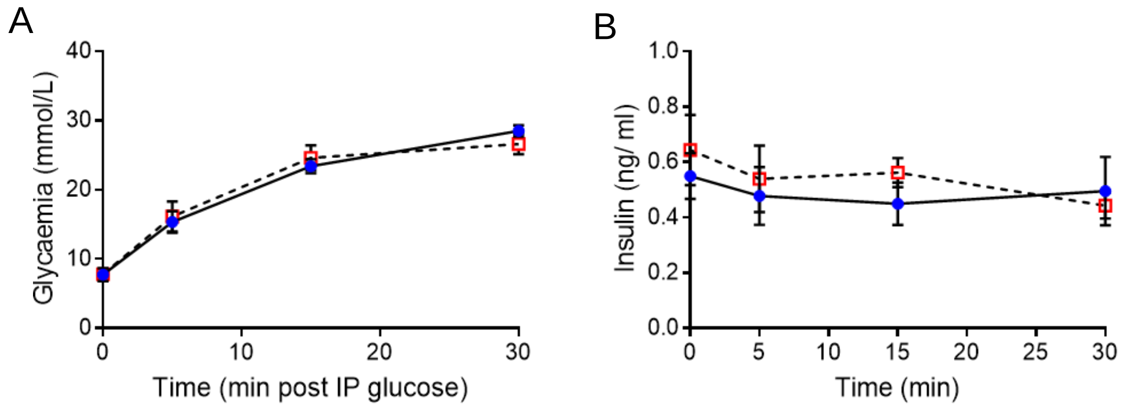
